# Macrophages from Holstein and Normande cows respond differently to *Escherichia coli* and *Streptococcus uberis*

**DOI:** 10.64898/2026.09.29.755538

**Authors:** Maryline Ferter, Christophe Gitton, Damaris F. Bamu, Marie Gaborit, Florence Gilbert, Pierre Germon

## Abstract

Macrophages, alongside mammary epithelial cells, are major contributors to the innate immune response triggered by bacterial invasion of the mammary gland. To investigate breed-specific immune mechanisms, we stimulated monocyte-derived macrophages from Holstein and Normande cattle with *Escherichia coli* and *Streptococcus uberis*, two major mastitis pathogens. We monitored bacterial survival using [méthode] and profiled gene expression via RNA-seq.

*E. coli* exhibited significantly higher survival 6 hours post-infection compared to *S. uberis* (p < 0.05). Moreover, *E. coli* induced a robust transcriptional response, with 4403 differentially expressed genes (FDR < 0.05, |log2FC| > 1), versus only 813 for *S. uberis*. Qualitative differences were also observed: genes such as CCL2 and CSF1 were uniquely induced by *S. uberis*. Additionally, macrophages from Holstein and Normande cattle showed distinct expression profiles: 185, 45, and 174 genes were differentially expressed at steady state or after *E. coli* or *S. uberis* stimulation, respectively. These genes were enriched in immune-related pathways.

Overall, our results demonstrate a stronger response to *E. coli*, although *S. uberis* specifically induces a subset of genes. The observed breed-specific differences may reflect underlying immune mechanisms that could influence disease susceptibility, warranting further functional validation.

## Introduction

Mastitis remains a major issue in dairy farming. Upon entry and recognition of bacterial pathogens by the host immune system, an inflammatory response is triggered which leads to recruitment of immune cells in the mammary gland [1].

The response of the mammary gland to infection by different pathogens is highly variable. In particular, while a strong and rapid inflammation is observed upon inoculation of *E. coli* in a mammary gland, the response to *S. uberis* inoculation is slower [2, 3]. This differential response is likely linked to different kinetics in the production of pro-inflammation mediators such as cytokines/chemokines or oxylipins and potentially to the involvement of different cell types in the initiation of the response [2–5].

In vitro data suggest that recognition of bacteria relies in particular on mammary epithelial cells and macrophages. While mammary epithelial cells have been shown to efficiently respond to stimulation by *E. coli* [6–9], conflicting data exist regarding their ability to respond to *S. uberis* infection [10, 11].

Available data rather suggest that milk macrophages are essential contributors to the onset of inflammation caused by *S. uberis* colonisation of the udder. While in vitro experiments suggest a low bactericidal capacity of milk macrophages toward *S. uberis*, macrophages, but not neutrophils, were shown to phagocytose *S. uberis* in experimentally infected mammary glands, suggesting that they play a key role in the defense against *S. uberis* infection [12–14].

Macrophages are indeed well equipped with different receptors that allow recognition of so-called Microbe-Associated Molecular Patterns carried by Gram-negative and Gram-positive pathogens [15]. A better understanding of the role macrophages play in the response to mammary gland infections requires a more detailed description of how bovine macrophages respond to mammary gland pathogens.

In order to address this question, we used blood monocytes derived macrophages (BMDM), as a surrogate for milk macrophages, to interrogate the response of macrophages to *S. uberis* infection and compared it to that following *E. coli* stimulation.

## Materials and methods

### Ethics Statement

Blood samples used in this study were collected with the approval from the ethics committee (agreements APAFiS #36413-2022040715544489_v3). Animal studies were compliant with all applicable provisions established by the European directive 2010/63/UE. The animals were submitted only to the introduction of needle in accordance with good veterinary practice and were not killed for the purpose of this study.

### Sample Collection

All blood samples were obtained from healthy Holstein cows housed at the Unité Expérimentale de Physiologie Animale de l’Orfrasière (INRAE, Nouzilly) or at the Unité Expérimentale du Pin (INRAE, Pin-au-Haras). For each cow, blood was collected into four 9 ml Vacutainer K2-EDTA tubes (BD Medical) by puncture in the jugular vein. The blood was kept at room temperature and processed within 2 hours of collection.

### Generation of Macrophages from Peripheral Blood Mononuclear cells

Whole blood was centrifuged at 1000g for 30 minutes at 20°C. Buffy coat was collected, diluted with two volumes of DPBS (without Mg, Ca). It was carefully layered onto 4mL of Percoll solution (d=1.077, GE Healthcare ref 17-0891-01). Tubes were centrifuged at 400g for 30 minutes, 20°C (Acceleration 8, Deceleration 2). The Peripheral Blood Mononuclear cells (PBMCs) were recovered, washed three times with DPBS (without Mg, Ca) containing 2mM EDTA and 2% Fetal Calf Serum (FCS), and resuspended 1 × 10^6^ cells/mL in Iscove’s Modified Dulbecco’s Medium-IMDM (Sigma, Aldrich) supplemented with 2mM glutamine, 10mM Hepes buffer, streptomycin (10mg/mL), penicillin (10,000 units) and amphotericin B (250ug/mL) before seeding in Sterilin Thermofischer plates. Plates were incubated at 37°C, with 5% CO_2_ for 2 hours in supplemented IMDM without serum to enable adherence of cells to the plates. Non-adherent cells were removed by performing three washes in DPBS then further incubated for 6 days. Medium change was done on the third day of incubation.

### Flow cytometry analyses

Macrophages obtained as described above were detached using accutase (Biolegend, Cat 423201, Lot B297842), adjusted to a final concentration of 10^5^ cells/mL before dispensing 50µL per well in a round-bottomed 96 well plate. The plate was centrifuged at 400g, 20°C, for 5 minutes. Cells were incubated with 100µL per well of Fixable Viability dye eFLuor 450 in PBS (ThermoFischer) for 20 minutes at 4°C. This was followed by two washes with FACs buffer (DBPS without Mg, Ca, 2mM EDTA+2% goat serum) addition of 50 µl final volume with the relevant antibodies diluted in FACs buffer. Isotype controls were performed with suitable antibodies (Table 1).

**Table 1:** Primary and Secondary Antibodies with Corresponding Isotypes.

| Marker | Antibody | Isotype | Manufacturer | Ref | Fluorophore |
| --- | --- | --- | --- | --- | --- |
| CD14 | Anti-CD14 | IgG2a | Biorad | MCA1568GA | A647 |
| MHCII | Anti-MHCII | IgG1 | VRMD | CAT82A |  |
| mIgG1 | Anti-mIgG1 |  | Biolegend | 406607 | PE |
| isotype CTL IgG1 |  | IgG1 | Biolegend | BLE401401 |  |
| CD11b | Anti-CD11b | IgG2b | Hybridoma |  |  |
| mIgG2b | Anti-IgG2b |  | BD Bioscience | 743175 | BV510 |
| isotype CTL IgG2b |  | IgG2b | Biolegend | BLE400301 |  |
| CD16 | Anti-CD16 | IgG2a | Biorad | MCA5665GA |  |
| mIgG2a | Anti-mIgG2a |  | Biorad | HCA037F | FITC |
| isotype CTL IgG2a |  | IgG2a | Biolegend | BLE400201 |  |

Flow cytometry was performed on a BD LSRFortessa X-20 cell analyzer (Becton-Dickinson). Data was analyzed using Kaluza analysis 2.1 (Beckman Coulter).

### Macrophage Stimulation with Interferon gamma

A day prior to stimulation, macrophages were detached using accutase (Biolegend, Cat 423201, Lot B297842) and seeded to a final concentration of 10^6^ cells/mL in Iscove medium – 10% FCS containing no antibiotics in a 48-well plate. Plates were incubated overnight at 37°C, 5% CO_2_ for cell attachment. On the day of infection, cells are washed once with DPBS (without Mg, Ca) before stimulation in Iscove-10% FCS medium with 200ng/mL bovine IFNγ (Biorad). Unstimulated cells served as negative controls. Plates were incubated for 24 hours at 37°C, 5% CO_2_ before analysis.

### Preparation of Bacterial cultures

The two strains of *S. uberis* were gifted by Dr. Florence Gilbert from INRAE facility. These strains MC1P4-1981ARG-2011-EL4 (referred to as MC1), of sequence type ST1076, and 2211AVD-EL17-2011 (referred to as 2211), of sequence type ST6, were isolated from cases of clinical and subclinical bovine mastitis respectively. The *E. coli* P4 is a prototypical mastitis isolate [16]. Aliquots of these bacteria were stored at -80°C in 10% DMSO up until the time for macrophage infection.

One day prior to macrophage infection, *S. uberis* was inoculated in BHI broths and incubated at 37°C with shaking 180rpm. A subculture was then made by streaking on Columbia agar with 5% Sheep Blood agar plate incubated at 37°C for 16-18 hours. Colonies were then scraped and diluted in 1.5mL DPBS (without Mg, Ca) and OD measured at 600nm.

*E. coli* was grown overnight in BHI broth at 37°C. Preliminary experiments allowed the estimation of CFUs from OD measurements at 600nm.

One OD_600_ corresponded to 1×10^9^ CFU/ml and 4.1×10^7^ CFU/ml for *S. uberis* and *E. coli* respectively.

### Macrophages infection with *S. uberis* and *E. coli*

Macrophages were generated as described above, harvested and adjusted to 1×10^6^ cells/mL in supplemented IMDM containing no antibiotics. Seeding was done in 48 well plates and incubated overnight for attachment at 37°C, 5% CO_2_. Prior to cell infection, cells were washed twice with DPBS (without Mg, Ca). Infection was done at a MOI of 1 (1 bacteria: 1 macrophage). CFUs were confirmed by performing 10-fold serial dilutions and doing overnight cultures at 37°C on TSA plates. Two hours post infection, supernatants were collected and stored at (-20°C) for further analysis. For survival assays, cells were washed thrice using DPBS (without Mg, Ca) and Iscove medium-10%FCS + 50µg/mL gentamicin was added to eliminate any extracellular bacteria. Thirty minutes later, in half of the wells, cells were washed and lysed with water adjusted to pH11 with NaOH and released bacteria were counted (intracellular bacteria at t=2h30). In the other wells, cells were washed thrice using DPBS (without Mg, Ca) and (Iscove medium -10%FCS) containing 50µg/mL gentamicin was added for 30 minutes to eliminate any extracellular bacteria before medium replacement with (Iscove medium-10%FCS) with 10 µg/mL gentamicin and further incubation at 37°C with 5% CO_2_ for 4 hours. Cells were then washed and lysed with water adjusted to pH11 with NaOH and released bacteria were counted (survived bacteria at t=6h). Bacterial survival was calculated at the percent of “survived bacteria” after the 6 hour incubation over the number of “intracellular bacteria at 2h30”.

### RNA Extraction and RT-qPCR

Six hour post-infection, RNA extraction was performed using the Nucleospin RNA kit (Macherey-Nagel). Cell culture medium was removed and macrophages were lysed with 200uL of RAI buffer before transferring the contents to a Nucleospin filter column and centrifuging at 11000g for 1 minute. The RNA binding conditions was adjusted by adding 200uL of 70% ethanol to the filtrate. This content was mixed thoroughly and then transferred to a new RNA spin column and centrifuged at 11000g for 30 seconds.

Next, 350uL of the membrane desalting buffer (MDB) was added to the column and centrifuged at 11000g, for 1 minute. 95uL of the DNase was introduced to the column and incubated at room temperature for 15 minutes. Three washes: once with 200uL buffer RAW2, twice with buffer RAW3 600uL, 250uL respectively was performed. RNA was finally eluted by using 60uL of RNase free water and centrifuged at 11000g for 1 minute.

### Illumina sequencing

Sequencing was performed by Helixio (France). Briefly, cDNA libraries were prepared from high quality RNA. Samples were tagged to allow subsequent identification, amplified by polymerase chain reaction (PCR) and quantified by quantitative PCR. Individual RNA-seq libraries were sequenced in duplicate at 150bp/sequence paired-end reads using an Illumina HiSeq 3000 sequencer (Illumina, USA).

### Differential Expression Analysis

Expression data analyses were performed using R studio (RStudio 2025.05.1). The edgeR package (version 4.7.3) was applied to identify statistically significant differentially-expressed genes (DEGs). The quantification file was filtered to keep only genes with at least one sample with a number of reads greater than 5. The dispersion estimation was performed using the GLMCommonDisp function. The glmFit function was used as the fit method and glmLRT function was used to determine DEGs following the inflammatory challenge. A threshold of abs(log_2_(FC)>1 and an FDR of 0.01 was set to select DEGs between unstimulated and *E. coli* or *S. uberis* stimulated macrophages. For breed differences analyses with a given stimulus, a threshold of abs(log_2_(FC)>0.6 and an adjusted p-value (FDR) of 0.05 were selected. DEG lists were analyzed using Ingenuity Pathway Analysis (v01-22-01) to investigate deregulated pathways.

## Results

### Bovine macrophages express CD14, CD11b, CD16 and MHC II

The method described above has been used in several studies to successfully purify bovine macrophages from PBMCs [13, 17]. To validate that cells obtained were indeed macrophages, we performed flow cytometry to measure the levels of expression of specific surface markers common with macrophages which are CD14, CD11b, MHC II and CD16 [18]. Flow cytometry analysis clearly showed our macrophages expressed CD14, CD11b, MHC II with few cells expressing CD16 (**Figure 1**).

**Figure 1:**
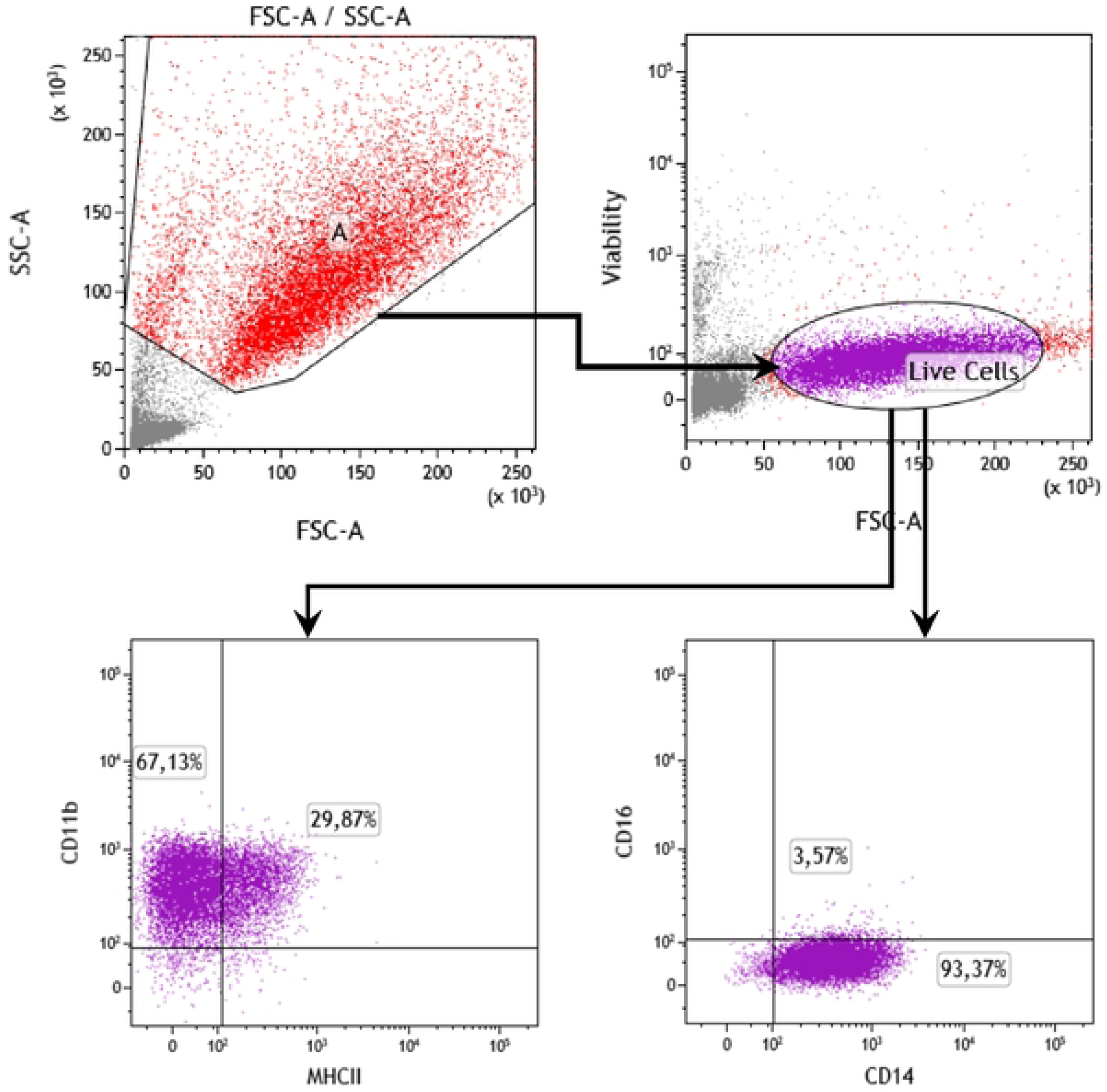
Cell surface markers expressed by bovine macrophages derived from PBMCs. Cells were washed with DPBS without Mg, Ca and detached with the aid of accutase. They were seeded 10^5^ cells/mL to 96-plate round bottom plate. We labelled them by incubating in 50uL of eFluorv450 viability dye, followed 50uL anti-mouse antibodies against CD11b, MHCII, CD16 and CD14. Flow cytometry was done using BD LSRFortessa X-20 cell analyzer.

### IFNγ modulates bovine macrophage surface markers

In order to check the functionality of the purified macrophages, since IFNγ had been associated with the activation of macrophages with enhanced MHC II expression [19], we stimulated them with recombinant bovine IFNγ. Following stimulation, cells were analysed for upregulation of cell surface markers. Increased expression of MHCII was detected with an increase from 29.08% to 65.66% for CD11b+/MHC II+ and a slight increase in CD16 expression, the percentage of CD14+/CD16+ increasing from 7.19% to 21.45% for (**Figure 2**).

**Figure 2:**
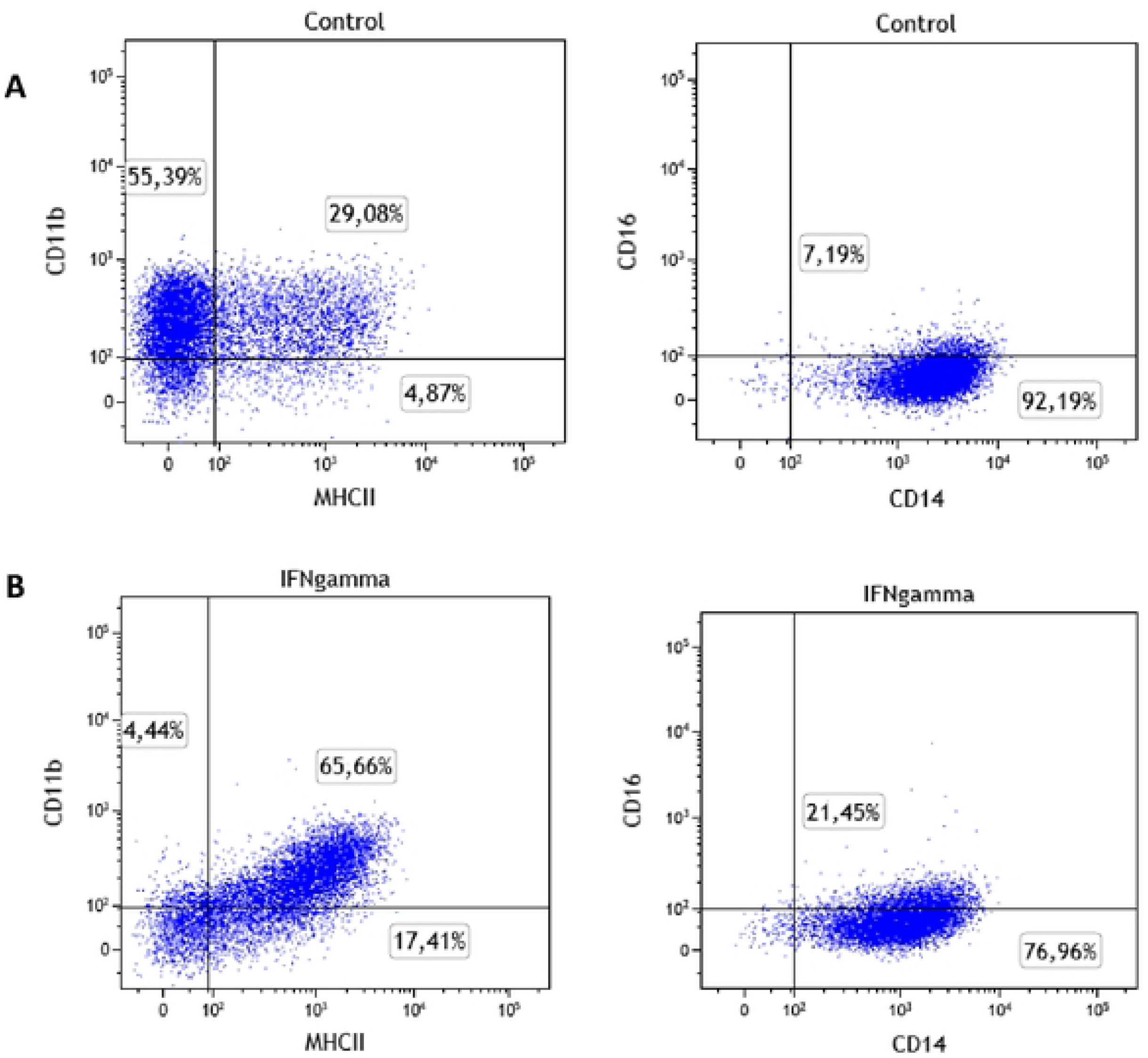
Effects of IFNγ on bovine macrophage surface markers. Detached macrophages were incubated overnight at 37°C, 5% CO_2_ prior to stimulation with 200ng/mL IFNγ (B) and incubation for 24 hours at 37°C, 5% CO_2_ or were left unstimulated (A). Cells were then detached and labelled with 50uL of viability dye, followed 50uL anti-mouse antibodies against CD11b, MHCII, CD16 and CD14.

### *S. uberis* is more susceptible to macrophage killing than *E. coli*

The above protocol was used to purify BMDM from eight Holstein and eight Normande cows. We first compared the ability of these purified macrophages to kill *E. coli* and *S. uberis*. The number of bacteria that survived killing by macrophages was significantly higher with *E. coli* P4 that with both *S. uberis* strains tested (**Figure 3**). All three strains were completely killed 24h post-infection. Holstein and Normande monocytes derived macrophages were equally able to kill both pathogens.

**Figure 3:**
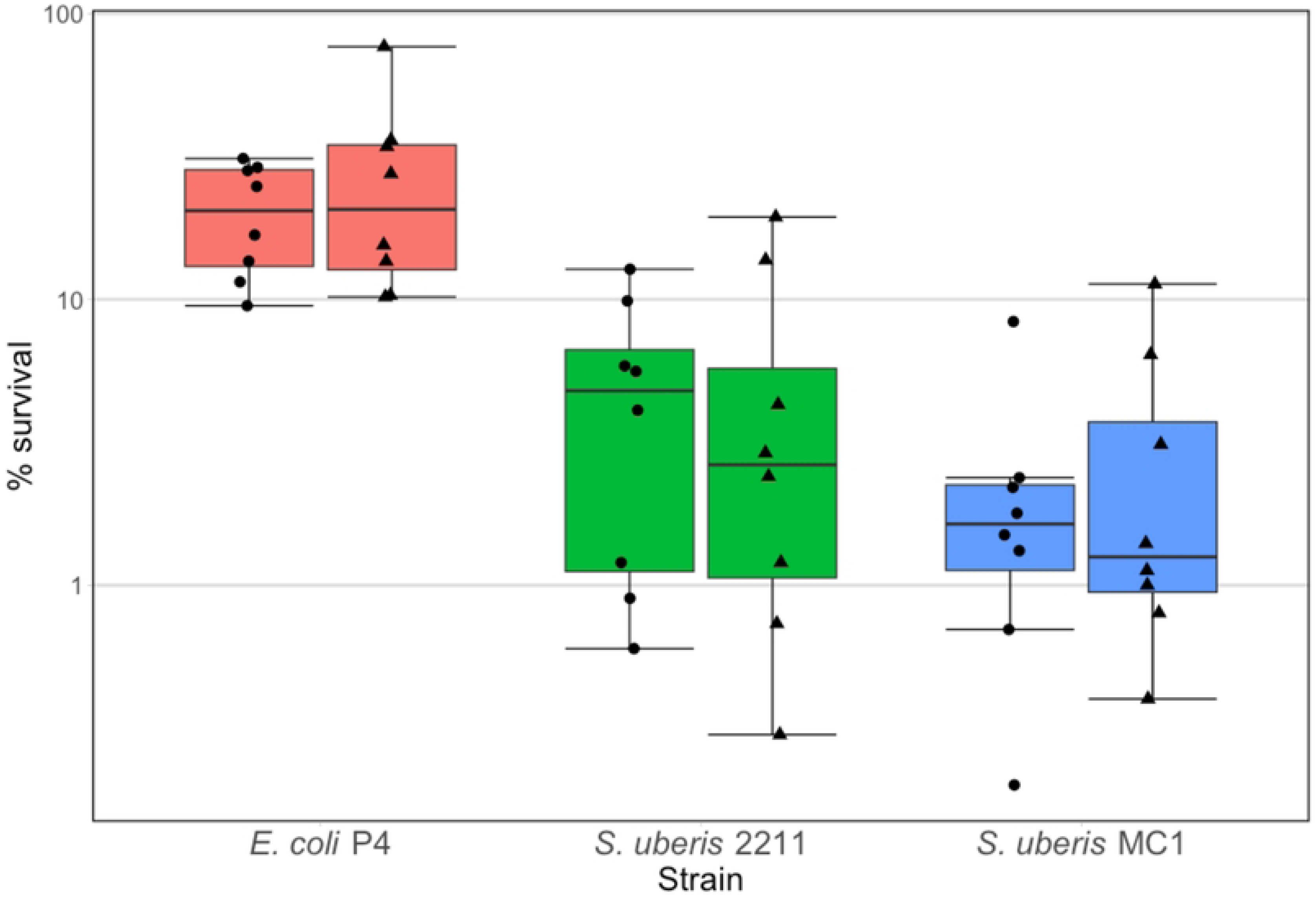
Survival of *E. coli* and *S. uberis* strains upon incubation with BMDM from Normande and Holstein cows. BMDM obtained from Holstein (●) and Normande (▲) cows were incubated at a MOI of 1 with either *E. coli* P4, *S. uberis* MC1 or *S. uberis* 2211 for 2 hours. Cells were then washed three times and medium with gentamicin (100 µg/ml) was added. Bacteria were recovered 4 hours later and counted. Survival is expressed as percent of the bacteria inside macrophages at t=2h.

### Response of BMDM to *E. coli* stimulation is stronger than that to *S. uberis*

We further explored the response of BMDM to *E. coli* and *S. uberis* by performing a transcriptomic analysis using RNA extracted from stimulated macrophages 6h p.i.. Because one RNA sample from Holstein cows was not of sufficient quality, we first analyzed the response of BMDM from Normande cows.

As expected, macrophages stimulated with *E. coli* responded very differently from *S. uberis* stimulated macrophages. Stimulation of Normande macrophages by *E. coli* led to 4403 differentially expressed genes (2299 up-regulated, 2104 down-regulated). Top 10 canonical pathways deregulated upon *E. coli* stimulation are listed in **Table 2**. Compared to the non-stimulated BMDM, stimulation by *S. uberis* led to deregulation of 813 genes, among which 643 were u-regulated and 170 down-regulated. Top 10 canonical pathways deregulated upon *S. uberis* stimulation are listed in **Table 3**.

**Table 2 :** List of top 10 deregulated pathways after stimulation of Normande BMDM with *E. coli*.

| Ingenuity Canonical Pathways | -log(p-value) | Ratio | z-score |
| --- | --- | --- | --- |
| Pathogen Induced Cytokine Storm Signaling Pathway | 1,67E01 | 4,34E-01 | 7,553 |
| Role of Hypercytokinemia/hyperchemokine-mia in the Pathogenesis<br>of Influenza | 1,62E01 | 7,19E-01 | 6,193 |
| Interleukin-4 and Interleukin-13 signaling | 1,52E01 | 5,82E-01 | 3,906 |
| Role of Pattern Recognition Receptors in Recognition of Bacteria<br>and Viruses | 1,42E01 | 5,33E-01 | 2,596 |
| IL-10 Signaling | 1,37E01 | 5,17E-01 | -0,465 |
| Th1 and Th2 Activation Pathway | 1,28E01 | 4,91E-01 | ND |
| Molecular Mechanisms of Cancer | 1,28E01 | 3,39E-01 | 2,625 |
| IL-17 Signaling | 1,27E01 | 4,97E-01 | 5,196 |
| Macrophage Alternative Activation Signaling Pathway | 1,24E01 | 4,76E-01 | 0,556 |
| NOD1/2 Signaling Pathway | 1,22E01 | 4,81E-01 | 3,989 |

**Table 3 :**
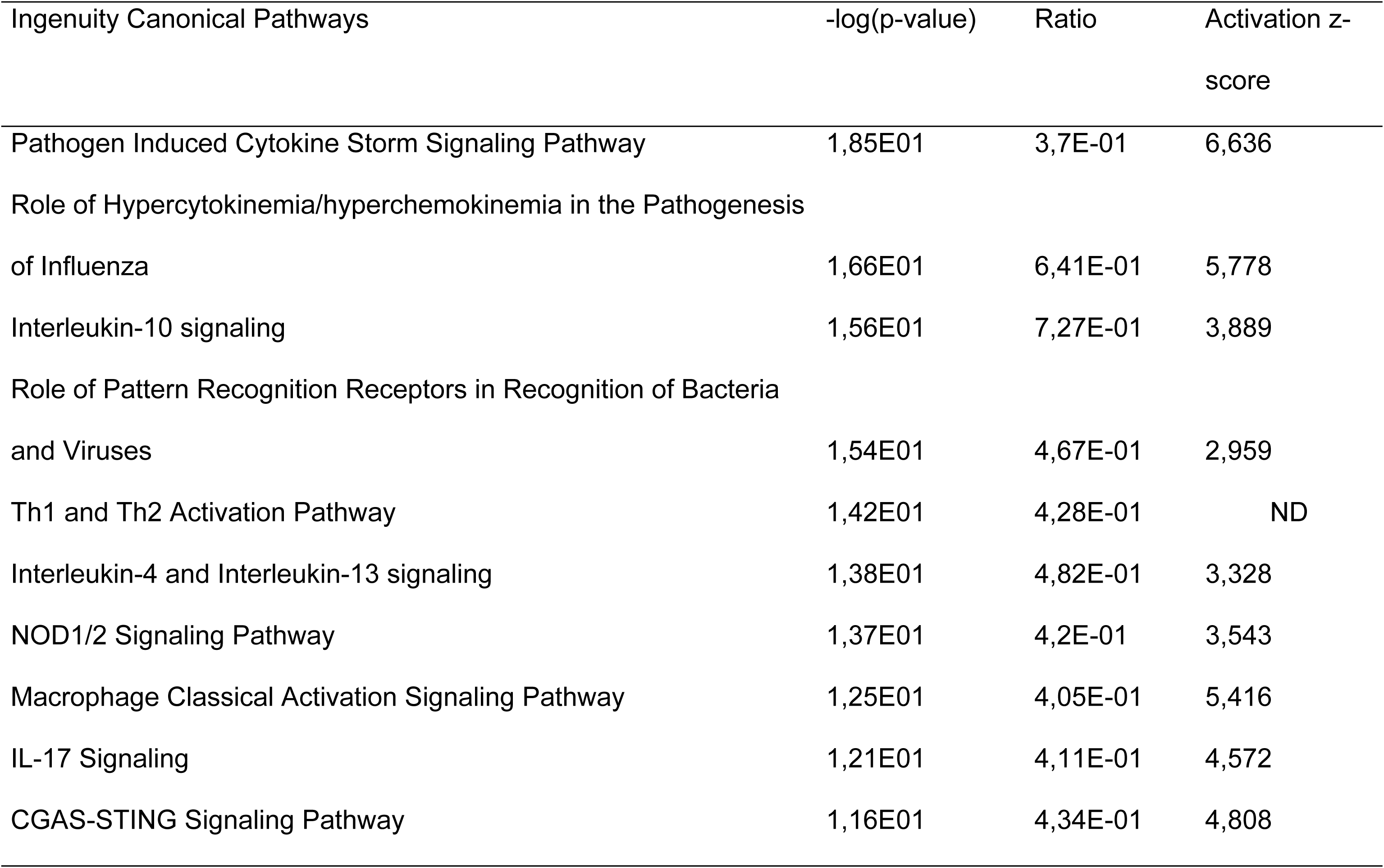
List of top 10 deregulated pathways after stimulation of Normande BMDM with *S. uberis*.

In both cases, not surprisingly, the « Pathogen Induced Cytokine Storm Signaling Pathway » was identified as the most deregulated pathway. Yet, a direct comparison of *E. coli*-stimulated versus *S. uberis* stimulated gene expression profiles identified 3231 DEG, of which 1695 were more expressed and 1536 were less expressed in *E. coli* stimulated samples. An interesting pattern was observed after detailed analysis of the « Pathogen Induced Cytokine Storm Signaling Pathway » which showed that contrary to most genes in this pathway, CCL2 was more expressed after stimulation with *S. uberis* (**Figure 4**). A closer investigation of genes that were up-regulated by *S. uberis* while down-regulated by *E. coli* stimulation identified a subset of 15 genes significantly induced (log2(FC) >1) after stimulation by *S. uberis* but not induced upon stimulation by *E. coli* (**Table 4**). Thus, the results of stimulation of macrophages by *S. uberis* is not only quantitatively weaker, it is also qualitatively different and suggests that some signaling pathways, in particular the one driving the expression of CCL2, are specifically activated by *S. uberis* but not, or to a lesser extent, by *E. coli*.

**Figure 4:**
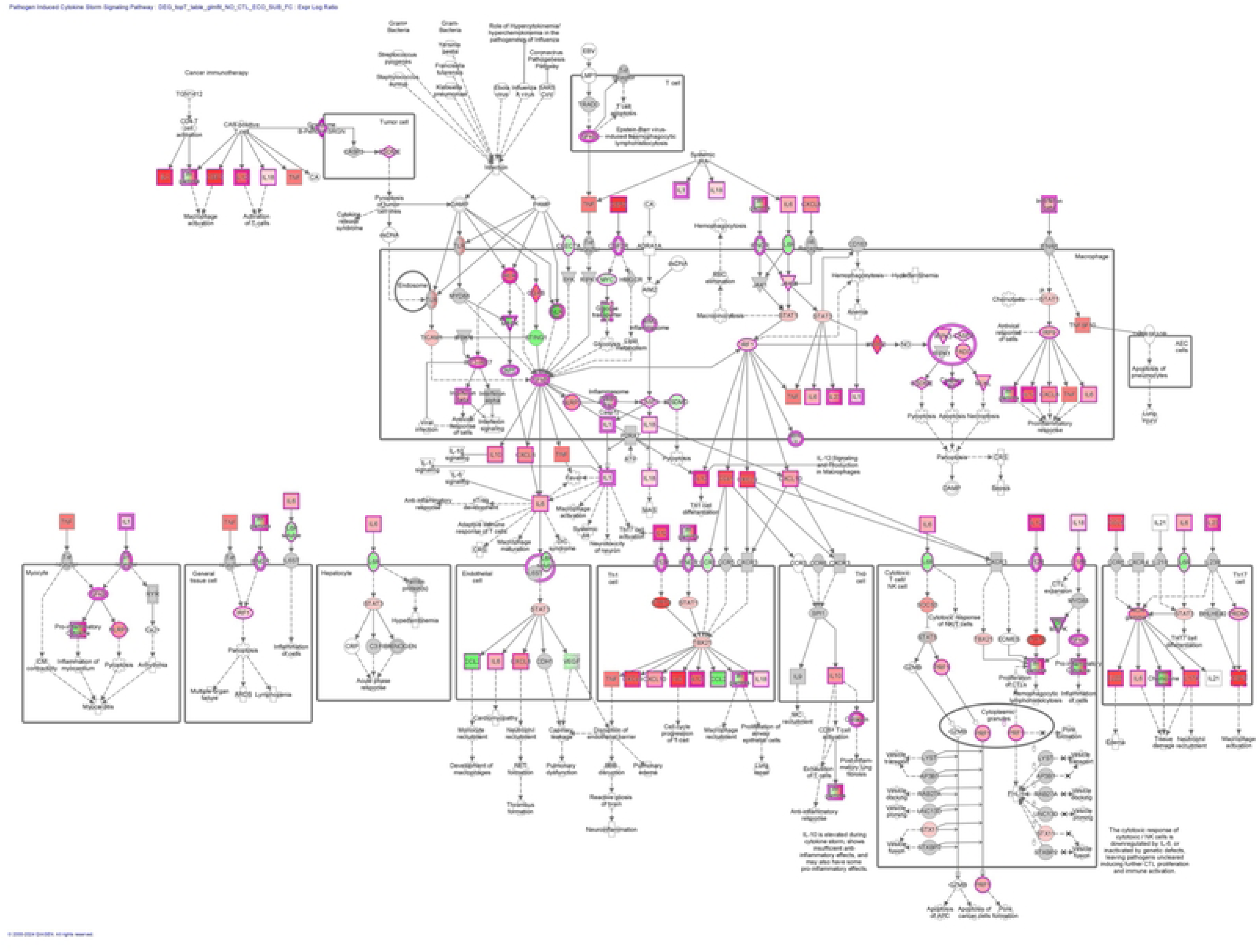
Graphical representation of the “Pathogen induced Cytokine Storm Signaling Pathway” genes log2FC in macrophages stimulated by *S. uberis* versus macrophages stimulated by *E. coli*. Red color gradient indicate over expression in *E. coli* stimulated macrophages while green color gradient indicates over expression in *S. uberis* stimulated macrophages.

**Table 4 :**
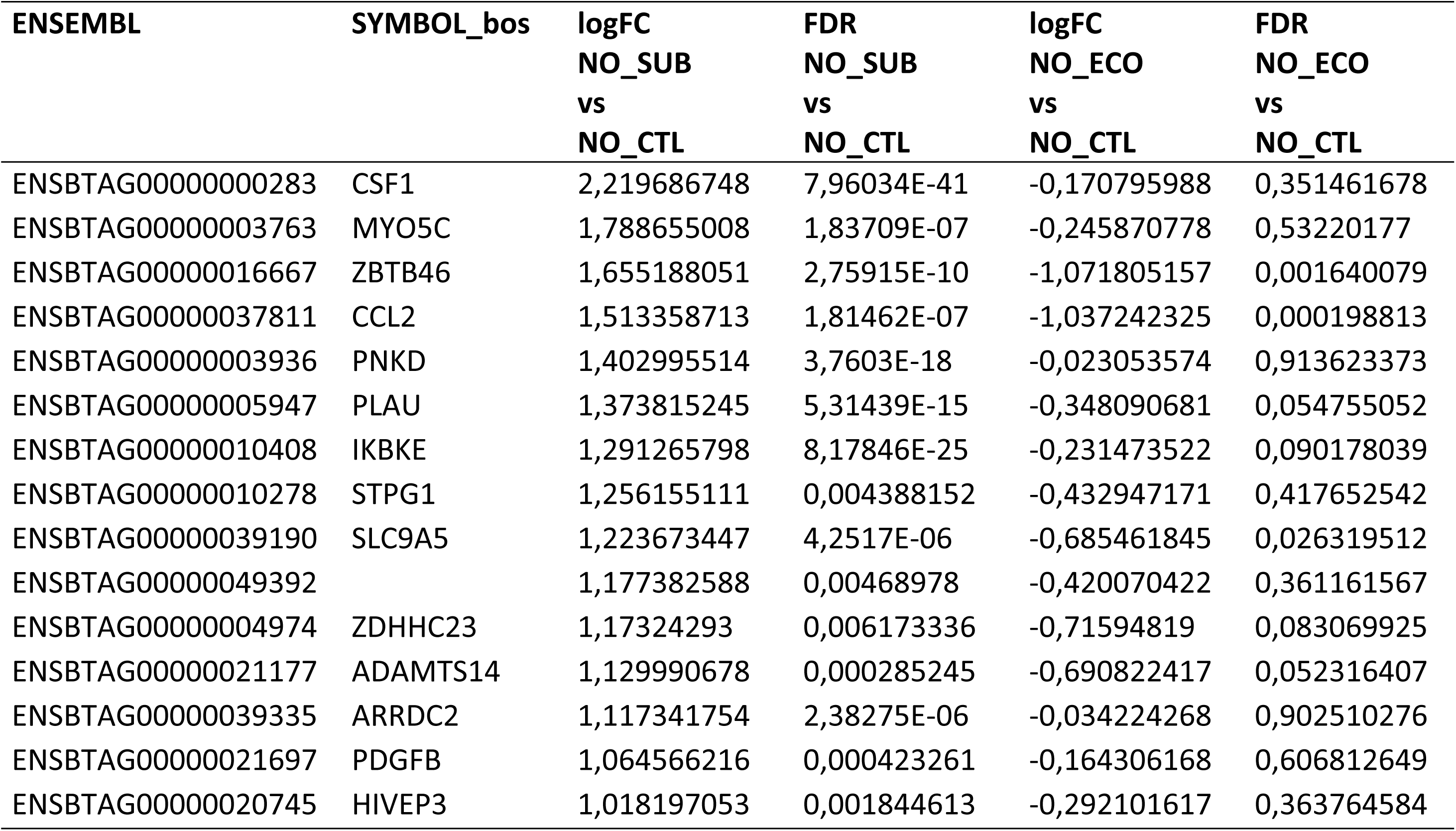
List of genes whose expression is induced upon stimulation of Normande BMDM with *S. uberis* but not after stimulation by *E. coli*.

When this analysis was performed on Holstein macrophages, despite missing one *S. uberis* stimulated sample because of RNA degradation, similar findings were obtained. In particular, the « Pathogen Induced Cytokine Storm Signaling Pathway » was the most deregulated pathway. RT-qPCR analyses, performed on all Holstein samples, including the one removed from RNAseq analyses due to RNA degradation, confirmed that CCL2 was also more expressed in *S. uberis* stimulated samples than in *E. coli* stimulated ones (data not shown).

### Response of Normande and Holstein BMDM to stimulation by *E. coli* or *S. uberis* reveals breed specific pathogen-responses

We then explored the possibility of differences between breeds. We thus separately compared the transcriptome of unstimulated BMDM from Holstein and Normande cows or upon stimulation with *E. coli* or *S. uberis* (**Figures 5A and 5B**). In the unstimulated situation, differences were observed with 185 DEG (151 up- and 34 down-regulated in Holstein macrophages). Upon stimulation, 45 (22 up- and 23 down-regulated in Holstein macrophages) and 174 DEG (119 up- and 55 down-regulated in Holstein macrophages) were detected between Holstein and Normande stimulated by *E. coli* or *S. uberis*, respectively. The complete list of DEG between breeds in each of the three conditions (unstimulated, *S. uberis* stimulation, *E. coli* stimulation) is provided in **Supplementary Table 1**. Investigation of the pathways differentially regulated between Normande and Holstein cows identified several pathways linked to immune response (**Table 5**). In particular, the four pathways « Pathogen Induced Cytokine Storm Signaling Pathway », « Interleukin-10 signaling », « IL-17 Signaling » and « Differential Regulation of Cytokine Production in Macrophages and T Helper Cells by IL-17A and IL-17F » were found significantly less activated in CTL and SUB conditions in Normande macrophages compared to Holstein macrophages (negative activation z-scores). Interestingly, upon stimulation with *E. coli*, which, as shown above, induces a stronger response than *S. uberis*, differences were less important and these four immune response related pathways were not identified as deregulated. A common set of 27 genes appeared to distinguish macrophages from Holstein from Normande cows in terms of response to pathogens (**Table 6**). Although the number of DEG was limited, four of these genes, *GBP4*, *TAP*, *MAPK12* and *LOC783604* (encoding a GBP4-like protein), are involved in host response to pathogen and belong to the Nod-signaling pathway.

**Table 5 :** List of IPA pathways differentially regulated between Holstein and Normande breeds in unstimulated conditions (CTL) or after stimulation by *S. uberis* (SUB) or *E. coli* (ECO)

| Ingenuity Canonical Pathways | P-value | Activation z-score | Percentage overlap |
| --- | --- | --- | --- |
| CTL |  |  |  |
| Pathogen Induced Cytokine Storm Signaling Pathway | 7.24e-20 | -4.80 | 7.83% |
| Interleukin-10 signaling | 1.97e-16 | -3.46 | 27.27% |
| Macrophage Classical Activation Signaling Pathway | 3.35e-15 | -4.00 | 10.24% |
| NOD1/2 Signaling Pathway | 9.87e-13 | -2.67 | 9.04% |
| Granulocyte Adhesion and Diapedesis | 1.97e-12 |  | 8.62% |
| IL-17 Signaling | 3.77e-12 | -3.74 | 9.33% |
| Differential Regulation of Cytokine Production in Macrophages and T Helper Cells by IL-17A and IL-17F | 1.55e-11 | -2.45 | 41.18% |
| Multiple Sclerosis Signaling Pathway | 2.25e-11 | -3.36 | 7.28% |
| Agranulocyte Adhesion and Diapedesis | 9.96e-11 |  | 7.33% |
| Irritable Bowel Syndrome Signaling Pathway | 2.08e-10 | -3.36 | 6.22% |
| SUB |  |  |  |
| Interleukin-10 signaling | 5.42e-11 | -2.83 | 18.18% |
| Class A/1 (Rhodopsin-like receptors) | 9.98e-10 | -3.61 | 4.89% |
| Granulocyte Adhesion and Diapedesis | 1.44e-9 |  | 6.32% |
| Pathogen Induced Cytokine Storm Signaling Pathway | 1.44e-8 | -3.32 | 3.92% |
| Agranulocyte Adhesion and Diapedesis | 5.08e-8 |  | 5.24% |
| IL-17 Signaling | 7.66e-8 | -3.00 | 6.00% |
| S100 Family Signaling Pathway | 2.18e-7 | -2.71 | 2.43% |
| G alpha (i) signalling events | 1.38e-6 | -3.00 | 4.27% |
| Differential Regulation of Cytokine Production in Macrophages and T Helper Cells by IL-17A and IL-17F | 1.48e-6 |  | 23.53% |
| Interleukin-4 and Interleukin-13 signaling | 1.65e-6 | -1.89 | 6.31% |
| ECO |  |  |  |
| Cholecystokinin/Gastrin-mediated Signaling | 2.23e-4 |  | 2.70% |
| CXCR4 Signaling | 6.76e-4 |  | 1.85% |
| BBSome Signaling Pathway | 8.38e-4 |  | 0.99% |
| Molecular Mechanisms of Cancer | 1.07e-3 | 0.00 | 0.66% |

**Table 6 :**
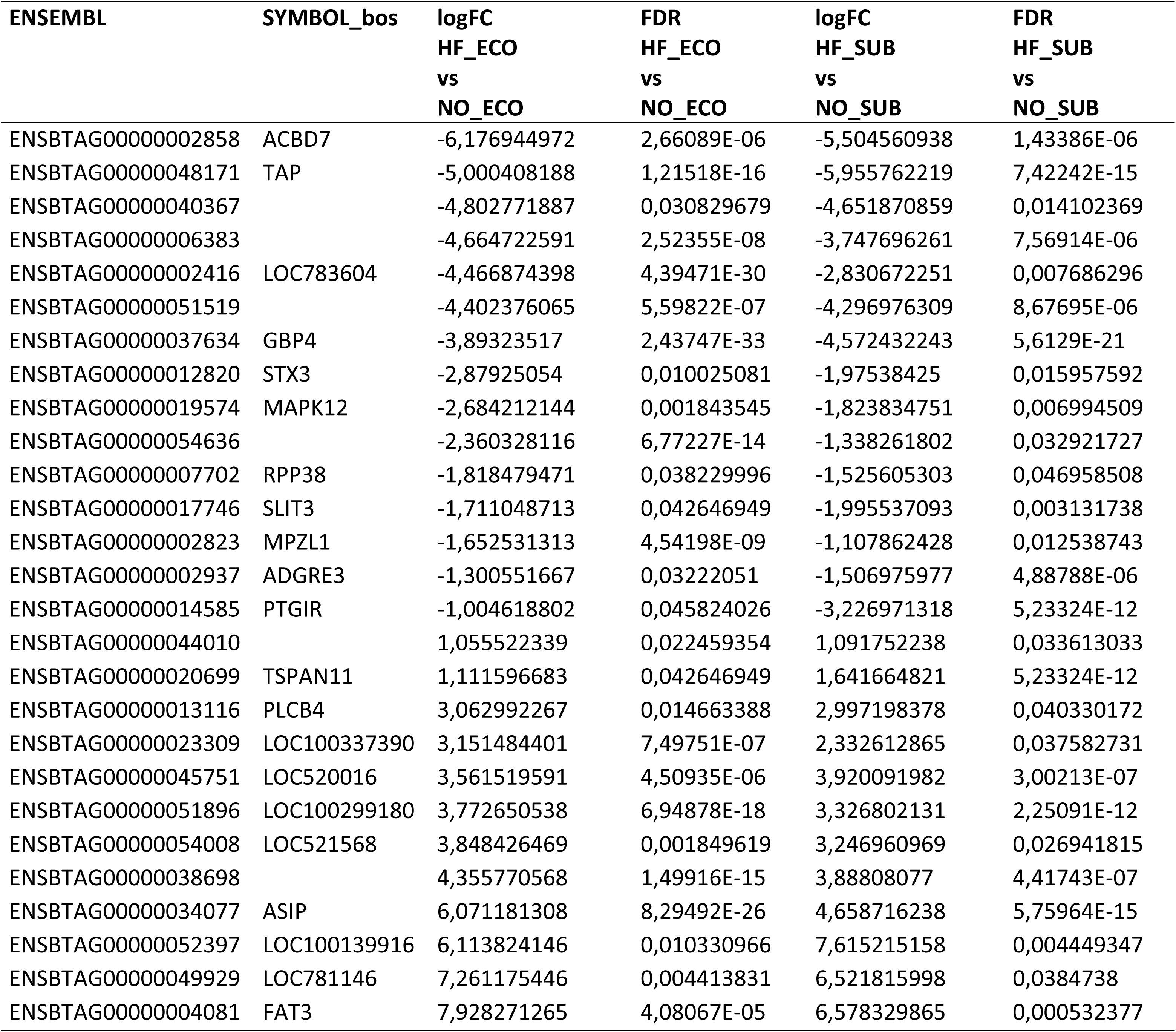
List of DEG between Normande and Holstein BMDM upon stimulation with *E. coli* or *S. uberis*.

**Figure 5:**
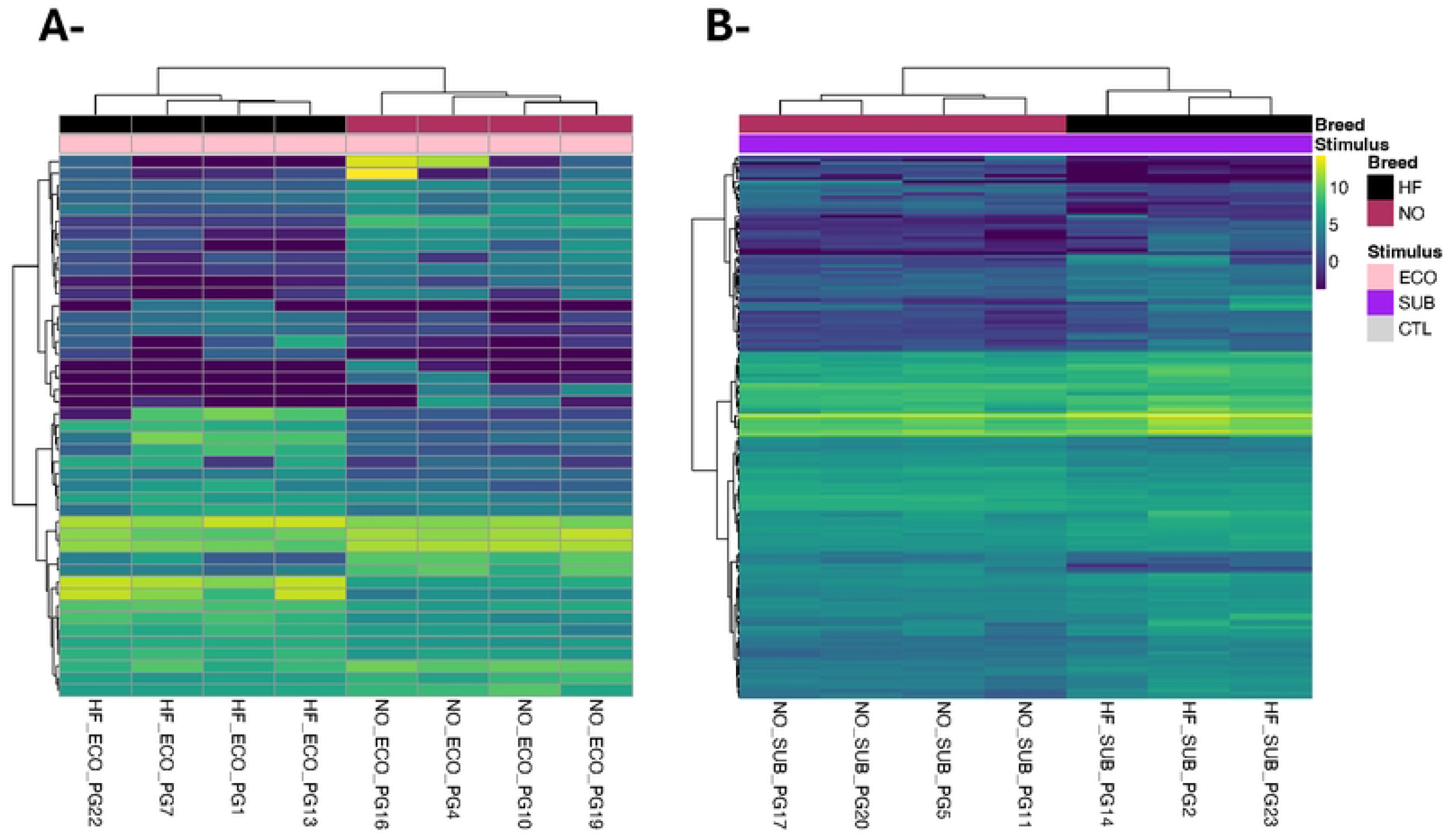
Heatmap and dot-plot representation of genes deregulated between Holstein and Normande cows after stimulation by *E. coli* of *S. uberis*. A-DEG between HF and NO macrophages after stimultation by *E. coli*. B-DEG between HF and NO macrophages after stimultation by *S. uberis*. C-dot-plot of log2FC values from comparison of HF vs NO macrophage stimulations by *S. uberis* and *E. coli*.

## Discussion

The objective of this study was to decipher the response of bovine macrophages to two main mammary gland pathogens, *E. coli* and *S. uberis*, and unravel potential breed-related differences in response to infection.

The relevance of using BMDM rather than milk macrophages is justified by previous studies which used BMDM to study interactions between macrophages and mastitis pathogens [20–22]. Most importantly, Denis *et al.* showed that, similar to dry secretion macrophages, BMDM were able to kill and respond by production of NO to infection by *S. uberis* while milk macrophages had lower response and were not able to kill *S. uberis* [13]. This is all the more relevant that interaction of *S. uberis* with dry secretion macrophages are likely to occur since a high percentage of *S. uberis* infections have been shown to be initiated during the dry period and experimental infection of dry glands has been demonstrated to be effective [23, 24] [25]. In addition, from a technical standpoint, using BMDM also allows for simultaneous stimulation of macrophages obtained from multiple cows and improve statistical power of the results. The functionality of the macrophages used in this report was first validated by verifying their response to IFNγ. In macrophages, interferon gamma (IFNγ) induces the expression of major histocompatibility complex (MHC) class II proteins that allows peptides coming from phagocytosed infectious agents to present to T cells [26]. Induction of MHCII was clearly detected in our macrophages. In addition, these macrophages were able to phagocytose and kill both *E. coli* and *S. uberis*.

Our results show that the response of bovine macrophages to *E. coli* leads to a higher number of DEG, either up- or down-regulated, compared to the response induced by *S. uberis*. This is not surprising given the very strong pro-inflammatory potential of the LPS exposed at the surface of *E. coli*. Yet, our results clearly identified specificities of the response to *S. uberis* where 15 genes were only up-regulated upon *S. uberis* stimulation, among which are found *CCL2* and *CSF1*.

In vivo, expression of *CCL2* is induced in the mammary gland tissue both after experimental infection by *E. coli* or *S. uberis* [27, 28]. In vitro data have shown that expression of *CCL2* was significantly increased in mammary epithelial cells stimulated by *E. coli* or LPS [29, 30]. Hence, it is possible that, upon *E. coli* infection, *CCL2* expression in vivo would be rather produced by MEC than by macrophages.

Nevertheless, the observation that *CCL2* expression is higher upon stimulation by *S. uberis* than with *E. coli* deserves further investigations since it suggests a species-specific induction pathway. A specific mechanism for the induction of *CCL2* by Streptococci has been observed in dendritic cells where the increased expression of *CCL2* was shown to be TLR2-independent while the induction of another chemokine, *CCL3*, was TLR2 and Myd88 dependent [31, 32]. Similarly, if TLR2 is also not involved in *S. uberis* recognition by macrophages, the contribution of particular lectin-type receptors should be investigated in more details: these receptors could in particular allow detection of some of the lectin-agglutination patterns possessed by different *S. uberis* isolates [33].

Among the other genes that show expression regulation upon infection similar to *CCL2*, *CSF1* is worth mentioning. The induction of *CSF1* by *S. uberis* but not *E. coli* could have functionnal consequences on macrophages activity since *CSF1* production is known to regulate differentiation of macrophages [34]. Whether this induction benefits *S. uberis* would require further investigation.

We also investigated in this study the differences of response between macrophages from Holstein and Normande cows. Upon stimulation by either pathogens or in the unstimulated condition, a number of genes were found differently regulated in Holstein and Normande macrophages. Ingenuity pathway analysis identified a number of pathways related to immune functions as less activated in Normande compared to Holstein macrophages.

Among the 27 genes differentially expressed in macrophages from Holstein from Normande cows, four of these genes, *GBP4*, *TAP*, *MAPK12* and *LOC783604* (encoding a GBP4-like protein), are involved in host response to pathogen and belong to the Nod-signaling pathway. *GBP4* is of particular interest: this gene encodes a GTPase belonging to a family of IFN*γ* inducible GTPases that are activated upon infection in murine IFN*γ* activated macrophages and contribute to the elimination of bacteria [37]. In cattle, GBP4 was significantly associated with bTB disease susceptibility and is significantly upregulated in blood from experimentally infected cattle [38, 39].

These differences raise the possibility that the two breeds show different susceptibilities to mastitis pathogens, and potentially to other pathogens. Yet, there are currently very few data regarding such breed differences, potentially because such comparisons are impeded by confounding effects such as differences in milk yield that is known to be negatively correlated to mastitis susceptibility [35, 36].

Regarding the limitations of this study, we investigated only one type of macrophages. Yet, macrophages represent a very diverse cell type, some having more pro-inflammatory activities while others are more efficient at wound healing for example. Although we tried to standardize our differentiation protocol, in particular by using the same batch of fetal calf serum for all experiments, we cannot exclude that different results could have been obtained with macrophages incubated with either CSF or GM-CSF to polarize them in different macrophage types.

In summary, this study provides a detailed analysis of macrophages response to infection by two major mastitis pathogens, *E. coli* and *S. uberis*. Although *E. coli* tends to trigger a stronger response from macrophages, the differences are not only quantitative since we were able to show that *S. uberis* is able to trigger an increased expression of some genes that are not activated by *E. coli*. Furthermore, our results, based on macrophages from two different bovine breeds, have identified a few genes that are differentially expressed between these two breeds; whether these differences translate into differences in susceptibility to disease remains to be verified.

## List of abbreviations

SCC: somatic cell count
DSCC: differential somatic cell count PMN: polymorphonuclear leukocytes
RT-qPCR: reverse transcription - quantitative polymerase chain reaction

## Declarations

### Ethics approval and consent to participate

Animal experiments were conducted in compliance with all applicable provisions established by the European directive 2010/63/UE. Blood sampling procedures were approved by local Ethics Committee (agreements APAFIS#29498-2021020410061759 and APAFIS#23901-20200203095344) . Our protocol did not interfere with farms breeding practices. All animals used in our study remained in the herd, without any space or food restriction.

### Consent for publication

Not applicable

### Availability of data and material

All data are available upon request. Full RNAseq data are available at the European Nucleotide Archive under the accession E-MTAB-17722.

### Authors’ contributions

This study was conceived by PG, MF and CG. MG provided resources. BFD, MF, CG and PG performed experiments. CG performed IPA analyses and prepared figures. PG designed, obtained funding, analyzed RNAseq data, prepared all figures and wrote the manuscript. All authors read and approved the manuscript before publication.

## Acknowledgment

Contribution to blood sampling of the INRAE experimental unit of Le Pin-au-Haras personnel is greatly acknowledged.

## References

1. Rainard P, Gilbert FB, Germon P. Immune defenses of the mammary gland epithelium of dairy ruminants. Frontiers in immunology. 2022;13:1031785. Epub 2022/11/08. doi: 10.3389/fimmu.2022.1031785. PubMed PMID: 36341445; PubMed Central PMCID: PMCPMC9634088.

2. Rambeaud M, Almeida RA, Pighetti GM, Oliver SP. Dynamics of leukocytes and cytokines during experimentally induced *Streptococcus uberis* mastitis. Vet Immunol Immunopathol. 2003;96(3-4):193–205. Epub 2003/11/01. doi: 10.1016/j.vetimm.2003.08.008. PubMed PMID: 14592732.

3. Bannerman DD, Paape MJ, Lee JW, Zhao X, Hope JC, Rainard P. Escherichia coli and *Staphylococcus aureus* elicit differential innate immune responses following intramammary infection. Clin Diagn Lab Immunol. 2004;11(3):463–72. Epub 2004/05/13. doi: 10.1128/CDLI.11.3.463-472.2004. PubMed PMID: 15138171; PubMed Central PMCID: PMCPMC404560.

4. Ryman VE, Pighetti GM, Lippolis JD, Gandy JC, Applegate CM, Sordillo LM. Quantification of bovine oxylipids during intramammary *Streptococcus uberis* infection. Prostaglandins Other Lipid Mediat. 2015;121(Pt B):207–17. Epub 2015/10/04. doi: 10.1016/j.prostaglandins.2015.09.006. PubMed PMID: 26432060.

5. Pezeshki A, Stordeur P, Wallemacq H, Schynts F, Stevens M, Boutet P, et al. Variation of inflammatory dynamics and mediators in primiparous cows after intramammary challenge with Escherichia coli. Vet Res. 2011;42(1):15. doi: 10.1186/1297-9716-42-15.

6. Strandberg Y, Gray C, Vuocolo T, Donaldson L, Broadway M, Tellam R. Lipopolysaccharide and lipoteichoic acid induce different innate immune responses in bovine mammary epithelial cells. Cytokine. 2005;31(1):72–86. Epub 2005/05/11. doi: 10.1016/j.cyto.2005.02.010. PubMed PMID: 15882946.

7. Yang W, Zerbe H, Petzl W, Brunner RM, Gunther J, Draing C, et al. Bovine TLR2 and TLR4 properly transduce signals from *Staphylococcus aureus* and *E. coli*, but *S. aureus* fails to both activate NF-kappaB in mammary epithelial cells and to quickly induce TNFalpha and interleukin-8 (CXCL8) expression in the udder. Mol Immunol. 2008;45(5):1385–97. Epub 2007/10/16. doi: 10.1016/j.molimm.2007.09.004. PubMed PMID: 17936907.

8. Gunther J, Esch K, Poschadel N, Petzl W, Zerbe H, Mitterhuemer S, et al. Comparative kinetics of Escherichia coli- and *Staphylococcus aureus*-specific activation of key immune pathways in mammary epithelial cells demonstrates that *S. aureus* elicits a delayed response dominated by interleukin-6 (IL-6) but not by IL-1A or tumor necrosis factor alpha. Infect Immun. 2011;79(2):695–707. Epub 2010/12/01. doi: 10.1128/IAI.01071-10. PubMed PMID: 21115717; PubMed Central PMCID: PMCPMC3028868.

9. Roussel P, Cunha P, Porcherie A, Petzl W, Gilbert FB, Riollet C, et al. Investigating the contribution of IL-17A and IL-17F to the host response during *Escherichia coli* mastitis. Vet Res. 2015;46(1):56. Epub 2015/06/13. doi: 10.1186/s13567-015-0201-4. PubMed PMID: 26062913; PubMed Central PMCID: PMCPMC4462179.

10. Gunther J, Czabanska A, Bauer I, Leigh JA, Holst O, Seyfert HM. *Streptococcus uberis* strains isolated from the bovine mammary gland evade immune recognition by mammary epithelial cells, but not of macrophages. Vet Res. 2016;47:13. Epub 2016/01/08. doi: 10.1186/s13567-015-0287-8. PubMed PMID: 26738804; PubMed Central PMCID: PMCPMC4704416.

11. Addis MF, Pisanu S, Marogna G, Cubeddu T, Pagnozzi D, Cacciotto C, et al. Production and release of antimicrobial and immune defense proteins by mammary epithelial cells following Streptococcus uberis infection of sheep. Infect Immun. 2013;81(9):3182–97. Epub 2013/06/19. doi: 10.1128/iai.00291-13. PubMed PMID: 23774600; PubMed Central PMCID: PMCPMC3754230.

12. Thomas LH, Haider W, Hill AW, Cook RS. Pathologic findings of experimentally induced *Streptococcus uberis* infection in the mammary gland of cows. Am J Vet Res. 1994;55(12):1723–8. Epub 1994/12/01. PubMed PMID: 7887517.

13. Denis M, Parlane NA, Lacy-Hulbert SJ, Summers EL, Buddle BM, Wedlock DN. Bactericidal activity of macrophages against *Streptococcus uberis* is different in mammary gland secretions of lactating and drying off cows. Vet Immunol Immunopathol. 2006;114(1-2):111–20. Epub 2006/09/05. doi: 10.1016/j.vetimm.2006.08.001. PubMed PMID: 16949677.

14. Pedersen LH, Aalbaek B, Rontved CM, Ingvartsen KL, Sorensen NS, Heegaard PM, et al. Early pathogenesis and inflammatory response in experimental bovine mastitis due to *Streptococcus uberis*. J Comp Pathol. 2003;128(2-3):156–64. Epub 2003/03/14. doi: 10.1053/jcpa.2002.0620. PubMed PMID: 12634093.

15. Werling D, Piercy J, Coffey TJ. Expression of TOLL-like receptors (TLR) by bovine antigen-presenting cells-potential role in pathogen discrimination? Vet Immunol Immunopathol. 2006;112(1-2):2–11. Epub 2006/05/17. doi: 10.1016/j.vetimm.2006.03.007. PubMed PMID: 16701904.

16. Bramley AJ, Faull WB, Young JL, Walton JR. Correspondence: Controlling coliform mastitis. Vet Rec. 1976;98(12):244. Epub 1976/03/20. doi: 10.1136/vr.98.12.244-a. PubMed PMID: 772945.

17. Denis M, Wedlock DN, Buddle BM. IFN-gamma enhances bovine macrophage responsiveness to *Mycobacterium bovis*: Impact on bacterial replication, cytokine release and macrophage apoptosis. Immunol Cell Biol. 2005;83(6):643–50. Epub 2005/11/04. doi: 10.1111/j.1440-1711.2005.01386.x. PubMed PMID: 16266317.

18. Hussen J, Schuberth HJ. Heterogeneity of Bovine Peripheral Blood Monocytes. Frontiers in immunology. 2017;8:1875. Epub 2018/01/10. doi: 10.3389/fimmu.2017.01875. PubMed PMID: 29312348; PubMed Central PMCID: PMCPMC5742132.

19. Su X, Yu Y, Zhong Y, Giannopoulou EG, Hu X, Liu H, et al. Interferon-gamma regulates cellular metabolism and mRNA translation to potentiate macrophage activation. Nat Immunol. 2015;16(8):838–49. Epub 2015/07/07. doi: 10.1038/ni.3205. PubMed PMID: 26147685; PubMed Central PMCID: PMCPMC4509841.

20. Tassi R, McNeilly TN, Sipka A, Zadoks RN. Correlation of hypothetical virulence traits of two Streptococcus uberis strains with the clinical manifestation of bovine mastitis. Vet Res. 2015;46:123. Epub 2015/10/27. doi: 10.1186/s13567-015-0268-y. PubMed PMID: 26497306; PubMed Central PMCID: PMCPMC4619225.

21. Farhat K, Sauter KS, Brcic M, Frey J, Ulmer AJ, Jungi TW. The response of HEK293 cells transfected with bovine TLR2 to established pathogen-associated molecular patterns and to bacteria causing mastitis in cattle. Vet Immunol Immunopathol. 2008;125(3-4):326–36. Epub 2008/07/16. doi: 10.1016/j.vetimm.2008.05.026. PubMed PMID: 18621422.

22. Lewandowska-Sabat AM, Kirsanova E, Klopp C, Solberg TR, Heringstad B, Østerås O, et al. Transcription Profiling of Monocyte-Derived Macrophages Infected In Vitro With Two Strains of Streptococcus agalactiae Reveals Candidate Pathways Affecting Subclinical Mastitis in Cattle. Front Genet. 2019;10:689. Epub 2019/08/17. doi: 10.3389/fgene.2019.00689. PubMed PMID: 31417606; PubMed Central PMCID: PMCPMC6681682.

23. Todhunter DA, Smith KL, Hogan JS. Environmental streptococcal intramammary infections of the bovine mammary gland. J Dairy Sci. 1995;78(11):2366–74. Epub 1995/11/01. doi: 10.3168/jds.S0022-0302(95)76864-3. PubMed PMID: 8747327.

24. McDonald JS, Anderson AJ. Experimental infection of bovine mammary glands with *Streptococcus uberis* during the nonlactating period. Am J Vet Res. 1981;42(3):465–7. Epub 1981/03/01. PubMed PMID: 7271011.

25. Hill AW. Pathogenicity of two strains of *Streptococcus uberis* infused into lactating and non-lactating bovine mammary glands. Res Vet Sci. 1988;45(3):400–4. Epub 1988/11/01. PubMed PMID: 3212290.

26. Tur J, Farrera C, Sánchez-Tilló E, Vico T, Guerrero-Gonzalez P, Fernandez-Elorduy A, et al. Induction of CIITA by IFN-γ in macrophages involves STAT1 activation by JAK and JNK. Immunobiology. 2021;226(5):152114. Epub 2021/07/26. doi: 10.1016/j.imbio.2021.152114. PubMed PMID: 34303919.

27. de Greeff A, Zadoks R, Ruuls L, Toussaint M, Nguyen TK, Downing A, et al. Early host response in the mammary gland after experimental *Streptococcus uberis* challenge in heifers. J Dairy Sci. 2013;96(6):3723–36. Epub 2013/04/17. doi: 10.3168/jds.2012-6320. PubMed PMID: 23587380.

28. Moyes KM, Sørensen P, Bionaz M. The Impact of Intramammary *Escherichia coli* Challenge on Liver and Mammary Transcriptome and Cross-Talk in Dairy Cows during Early Lactation Using RNAseq. PLOS ONE. 2016;11(6):e0157480. doi: 10.1371/journal.pone.0157480.

29. Gunther J, Koczan D, Yang W, Nurnberg G, Repsilber D, Schuberth HJ, et al. Assessment of the immune capacity of mammary epithelial cells: comparison with mammary tissue after challenge with *Escherichia coli*. Vet Res. 2009;40(4):31. Epub 2009/03/27. doi: 10.1051/vetres/2009014. PubMed PMID: 19321125; PubMed Central PMCID: PMCPMC2695127.

30. Vedrine M, Gilbert FB, Maman S, Klopp C, Gitton C, Rainard P, et al. Soluble CD14 produced by bovine mammary epithelial cells modulates their response to full length LPS. Vet Res. 2024;55(1):76. Epub 2024/06/13. doi: 10.1186/s13567-024-01329-3. PubMed PMID: 38867337; PubMed Central PMCID: PMCPMC11170775.

31. Goyette-Desjardins G, Auger JP, Dolbec D, Vinogradov E, Okura M, Takamatsu D, et al. Comparative Study of Immunogenic Properties of Purified Capsular Polysaccharides from *Streptococcus suis* Serotypes 3, 7, 8, and 9: the Serotype 3 Polysaccharide Induces an Opsonizing IgG Response. Infect Immun. 2020;88(10). Epub 2020/08/05. doi: 10.1128/IAI.00377-20. PubMed PMID: 32747605; PubMed Central PMCID: PMCPMC7504959.

32. Calzas C, Goyette-Desjardins G, Lemire P, Gagnon F, Lachance C, Van Calsteren MR, et al. Group B *Streptococcus* and *Streptococcus suis* capsular polysaccharides induce chemokine production by dendritic cells via Toll-like receptor 2- and MyD88-dependent and -independent pathways. Infect Immun. 2013;81(9):3106–18. Epub 2013/06/19. doi: 10.1128/IAI.00113-13. PubMed PMID: 23774593; PubMed Central PMCID: PMCPMC3754219.

33. Almeida RA, Oliver SP. Growth curve, capsule expression and characterization of the capsular material of selected strains of *Streptococcus uberis*. Zentralbl Veterinarmed B. 1993;40(9-10):697–706. Epub 1993/12/01. doi: 10.1111/j.1439-0450.1993.tb00194.x. PubMed PMID: 8128804.

34. Sehgal A, Irvine KM, Hume DA. Functions of macrophage colony-stimulating factor (CSF1) in development, homeostasis, and tissue repair. Semin Immunol. 2021;54:101509. Epub 2021/11/08. doi: 10.1016/j.smim.2021.101509. PubMed PMID: 34742624.

35. Heringstad B, Klemetsdal G, Steine T. Selection responses for clinical mastitis and protein yield in two Norwegian dairy cattle selection experiments. J Dairy Sci. 2003;86(9):2990–9. Epub 2003/09/26. doi: 10.3168/jds.S0022-0302(03)73897-1. PubMed PMID: 14507036.

36. Govignon-Gion A, Dassonneville R, Baloche G, Ducrocq V. Multiple trait genetic evaluation of clinical mastitis in three dairy cattle breeds. Animal. 2016;10(4):558–65. Epub 2015/11/26. doi: 10.1017/S1751731115002529. PubMed PMID: 26592099.

37. Tretina K, Park E-S, Maminska A, MacMicking JD. Interferon-induced guanylate-binding proteins: Guardians of host defense in health and disease. J Exp Med. 2019;216(3):482–500. doi: 10.1084/jem.20182031.

38. O’Grady JF, McHugo GP, Ward JA, Hall TJ, Faherty O’Donnell SL, Correia CN, et al. Integrative genomics sheds light on the immunogenetics of tuberculosis in cattle. Commun Biol. 2025;8(1):479. Epub 2025/03/25. doi: 10.1038/s42003-025-07846-x. PubMed PMID: 40128580; PubMed Central PMCID: PMCPMC11933339.

39. Correia CN, McHugo GP, Browne JA, McLoughlin KE, Nalpas NC, Magee DA, et al. High-resolution transcriptomics of bovine purified protein derivative-stimulated peripheral blood from cattle infected with Mycobacterium bovis across an experimental time course. Tuberculosis. 2022;136:102235. doi: 10.1016/j.tube.2022.102235.

